# A simple, flexible and high-efficiency western blot analysis for age-related human induced neurons

**DOI:** 10.1101/2023.01.30.526150

**Authors:** Yan-Fei Shen, Ming-Jie Liu, Zhu Long, Xiaobang Shi, Meng-Lu Liu

## Abstract

High-throughput western blot (WB) analysis of small and precious samples, such as various age-related subtype-specific human induced neurons (hiNs), confers the ability to obtain more consistent, comparable, and informative data from materials with extremely limited availability. In this study, p-toluenesulphonic acid (PTSA), an odorless tissue fixative, was used to inactivate HRP for developing a high-throughput WB method. PTSA-treated blots showed fast and efficient inactivation of HRP without detectable protein loss and epitope damage. With a brief PTSA-treatment before every next probing, 10 proteins of dopaminergic hiNs could be sequentially, sensitively, and specifically detected in a blot. These WB data proved the age-associated and neuron-specific features of hiNs and further revealed a sharp reduction of two Parkinson’s disease-associated proteins, UCHL1 and GAP43, in the normal aging dopaminergic neurons. Together, this study developed a unique and high-efficiency WB analysis and pinpointed its special value for capturing robust useful data from limited, precious samples.

**GRAPHICAL ABSTRACT:** 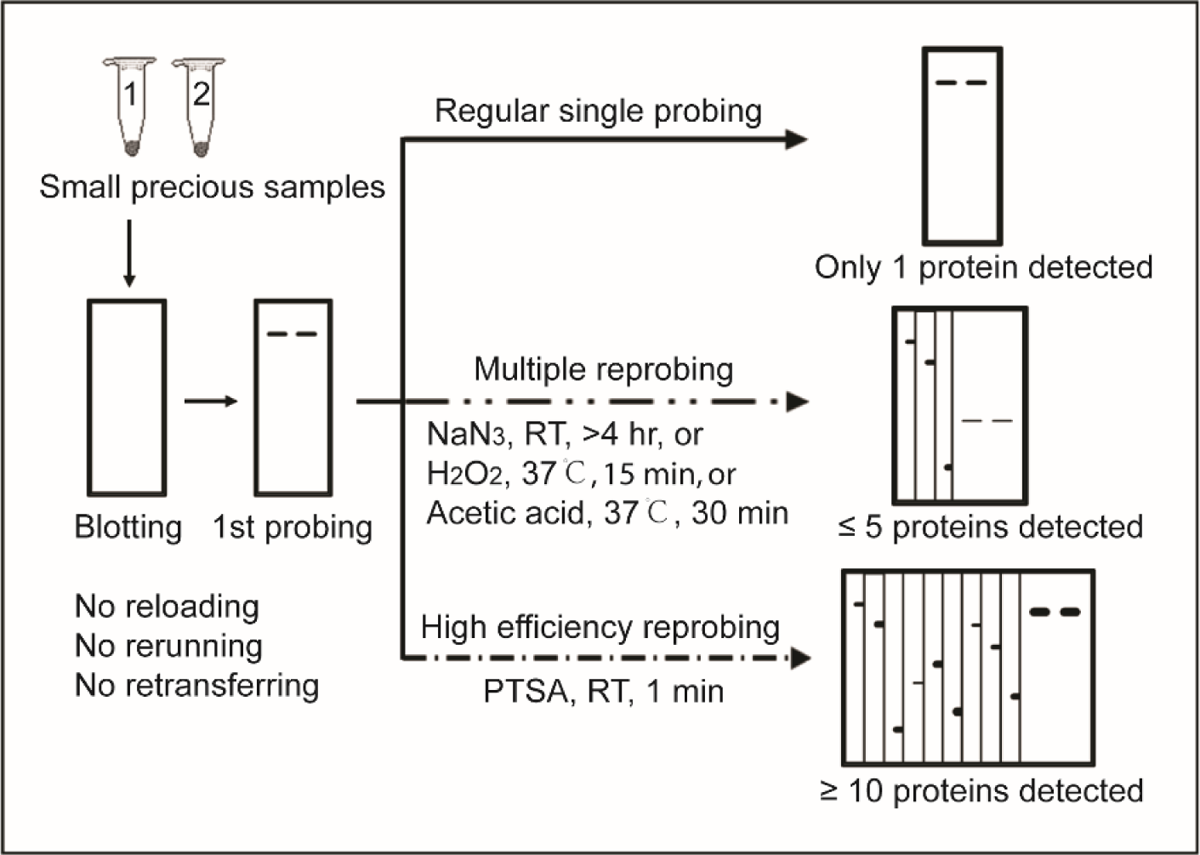

**HIGHLIGHTS:**

✓ P-toluenesulphonic acid (PTSA) quickly and fully deactivated HRP on immunoblots.
✓ PTSA was an odorless, non-volatile, low cost, and user-friendly HRP inactivator.
✓ PTSA allowed high-efficiency WB analysis to save small precious samples and time.
✓ 10 proteins were detected in a single blot of age-relevant human induced neurons.
✓ UCHL1 and GAP43 sharp decline occurred in aging human induced dopaminergic neurons.

## INTRODUCTION

Human induced neurons (hiNs) directly converted from skin fibroblasts do not pass through a proliferative progenitor state and keep the age-relevant information and disease-specific features of their parental cells [1-6]. In contrast, induced pluripotent stem cells (iPSCs)-derived neurons have been reset to a fetal status absence of aging- and/or disease-associated gene expression profiles [4-7]. Therefore, hiNs appear to be more proper disease models for screening therapeutics, as well as for understanding the age-related Alzheimer’s disease (AD), Parkinson’s disease (PD), and other neurodegenerative diseases [7-12]. The age-related and disease-specific gene expression patterns of hiNs have been mainly revealed through bioinformatical and immunocytochemical approaches [4-6]. However, due to the limited availability of highly pure subtype-specific hiNs [12], it remains difficult to use regular western blot (WB) analysis to convincingly validate the transcriptional and cellular findings, and to catch more physiological, pathological, and pharmacological insights from the aged and/or diseased cells. Thus, we are prompted to develop a proper method for quickly and efficiently reprobing an immunoblot loaded with much less proteins of hiNs.

In general, a variety of proteins blotted on the membrane could be labelled by their specific antibody and then the corresponding HRP conjugated secondary antibody, followed by an enhanced chemiluminescence (ECL) detection. This potentially provides the ability of multiple reprobing for one single blot to sequentially detect different proteins, instead of rerunning several gels and blots, in order to save precious sample and time. Stripping and HRP inactivation are two major methods for reprobing blots. Although diverse stripping reagents have been commercially available, the stripping procedures are usually very lengthy, mostly accompanied with strong odor, and often cause unavoidable antigen loss [13, 14]. Alternatively, HRP inactivation is easier to handle and thus more favorable for the reprobing purpose [15-18]. It is reasonable to speculate that a fast and efficient HRP-inactivating method would allow to multiply and quickly reprobe the blot for diverse proteins, leading to a high efficiency assay for protein samples with limited resources.

Several reagents have been applied to treat the blot for HRP inactivation. They include chromogenic substrate such as 3.3-diaminobenzidine [15], sodium azide [16], H_2_O_2_ [17], and acetic acid (AA) [18]. However, each solution has its own drawbacks. Chromogenic substrates are mutagenic and form insoluble precipitation that might affect further detection [15]. Sodium azide (NaN_3_) is a toxic, mutagenic, and explosive reagent, and inactivates HRP inefficiently, requiring long incubation time (≥ 4 hr) [16]. Treatment with high concentration H_2_O_2_ (appropriate condition: 30%, 30 min, 37℃) is an alternative way to inactivate HRP; however, high concentration H_2_O_2_ may induce an unpredictable effect on epitope recognition due to oxidation of susceptible epitopes [19]. Therefore, H_2_O_2_-treated blots may need to be re-blocked for further probing. Additionally, H_2_O_2_ has a slightly irritating odor, and appears to be an animal carcinogen and mutagen [19]. Under a pre-optimized, case-sensitive condition, AA is also able to inactivate HRP, but it may need further optimization on a case-by-case basis, and should be used with caution because it is a strong eye, skin, and mucous membrane irritant, with high volatility under the applicable condition [18].

The aqueous solution of p-toluenesulphonic acid (PTSA) has been used as an alternative fixative for the central nervous system [20]. PTSA-fixed cells and tissues preserved good morphology and histochemical properties [20]. As a non-volatile and non-oxidizing strong organic acid, PTSA is odorless, highly soluble, easy to handle, and extremely low-cost. In this study, we found that 0.05-0.5 M PTSA treatment of the blot for only 1 min at room temperature (RT) could quickly and fully inactivate HRP without any undesirable effect on the subsequent immunodetection. This allowed PTSA to be used as an excellent user-friendly HRP-inactivating reagent for a novel high-throughput WB analysis, with great value for studying precious highly pure hiNs.

## MATERIAL AND METHODS

### Antibodies and reagents

The primary and secondary antibodies used in this study were listed in Table 1. Nitrocellulose (NC) membrane (YA1711) was ordered from Solarbio, PRC. [PVDF](https://omnexus.specialchem.com/selection-guide/polyvinylidene-fluoride-pvdf-plastic) membrane (ISEQ00010) was from Milipore, USA. Acetic acid ([A116172](https://www.aladdin-e.com/zh_cn/catalogsearch/result/?q=64-19-7)) and p-toluenesulfonic acid monohydrate (T305333) were from Aladdin, PRC. Enhanced chemiluminescent kit (P0018AS) and other reagents, unless otherwise indicated, were purchased from Beyotime, PRC.

**Table 1.** Primary and secondary antibodies used in this study.

| Primary Antibody | Host | Identifier | Supplier | Dilution |
| --- | --- | --- | --- | --- |
| β-ACTIN | Mouse | 66009-1 | ProteinTech, USA | 1:10K |
| LGALS3 (Gal3) | Mouse | M00621-3 | Boster, USA | 1:10K |
| TUJ1 | Mouse | A5107 | Selleck, USA | 1:10K |
| ChAT | Rabbit | A5303 | Selleck, USA | 1:10K |
| GAPDH | Rabbit | 10094-T52 | Sino Biological, PRC | 1:10K |
| PBR/TSPO | Rabbit | IPB5348 | Baijia Biotech, PRC | 1:5K |
| Synapsin1 (SYN1) | Rabbit | D12G5 | CST, USA | 1:5K |
| LAMIN A/C | Rabbit | A5101 | Selleck, USA | 1:5K |
| UCHL1 (PGP9.5) | Mouse | M01018-6 | Boster, USA | 1:2K |
| LAMIN B1 | Mouse | 66095-1 | ProteinTech, USA | 1:5K |
| GAP43 | Rabbit | 16971-1 | ProteinTech, USA | 1:2K |
| BCL-2 | Rat | ab241548 | Abcam, USA | 1:5K |
| DAT | Rabbit | 22524-1-AP | ProteinTech, USA | 1:2K |
| Tyrosine hydroxylase (TH) | Mouse | 66334-1-Ig | ProteinTech, USA | 1:2K |
| **Secondary Antibody** |  |  |  |  |
| Anti-mouse IgG-HRP | Donkey | 715-035-151 | Jackson, USA | 1:10K |
| Anti-rabbit IgG-HRP | Donkey | 711-035-152 | Jackson, USA | 1:10K |
| Anti-rat IgG-HRP | Donkey | 712-035-150 | Jackson, USA | 1:10K |

### Cell resource and lysis

The highly unique resource materials of fibroblast-converted human induced basal forebrain cholinergic neurons (ECT-BFCN601) and human induced dopaminergic neurons (ECT-DN101 and ECT-DN601), as well as their parental fibroblasts were from frozen samples in Ecyton Biotech, PRC. The young DN101 was directly generated from human new born fibroblasts (#2310, Sciencell, USA); and the old DN601 or BFCN601 was separately generated from aging fibroblasts (66 YR, AG08517, Coriell, USA) using the commercially available trans-differentiation kits corresponding to each subtype of neurons (ECT-DNR01 and ECT-BFCNR01, Ecyton, PRC). The highly pure neurons or fibroblasts were directly lysed for WB assay, using a lysis buffer (P0013, Beyotime, PRC) containing protease inhibitors (B14001, Bimake, USA) and phosphatase inhibitors (B15001, Bimake, USA).

### Dot blot analysis

All antibodies were diluted in the phosphate buffered saline containing 0.1% Tween-20 (v/v) (PBST), 5% BSA, and 0.2% ProClin 300 (48912-U, Sigma, USA). For dot blotting assay, donkey anti-mouse IgG-HRP secondary antibody was sequentially diluted into 5 concentrations as indicated. Then, 1 μl of each dilution was spotted onto the NC membrane in quintuplicate or sextuplicate and dried at 37℃ for 30 min. After briefly rinsing with PBST, the initial HRP activity of all spots was detected using a regular ECL method. Then, the dot blots were cut into several equal parts, followed by treatment with either AA or PTSA. To completely remove any residue AA or PTSA, the blots were washed three times in PBST each for 10 min. Fresh ECL substrates were further applied to detect any remaining HRP activity on the acid-treated spots. All steps of rinse and acid treatment were performed at RT with constant shaking.

### Western blot analysis

WB assay was performed at RT according to a previous protocol with slight modification [10]. Briefly, denatured protein samples (10 μg young fibroblast lysate or 1 μg neuron lysate per lane) were loaded onto 12.5% (w/v) sodium dodecyl sulfate-polyacrylamide gels (SDS-PAGE) for electrophoresis, and then transferred onto PVDF membrane. After blocking with 5% non-fat milk in PBST for 2 hr, the blots were incubated with primary antibodies for another 2 hr, and then washed three times in PBST with constant shaking for 10 min. Afterward, the blots were incubated with HRP-conjugated secondary antibodies for 1 hr, and then washed three times again, followed by chemiluminescent detection, PTSA treatment, and reprobing.

### PTSA treatment

PTSA was dissolved in highly pure water to make a 0.5 M stock solution. NC or PVDF membrane blots were washed with PBST or treated with acid solution (either 10% AA or 0.005-0.5 M PTSA) in plastic boxes at RT with gentle constant shaking. For consistent results, excess solution (2 ml for 1 cm^2^) should be used to treat the blots. After every acid treatment, the blots were thoroughly rinsed for three times in PBST with constant shaking unless otherwise noted. Except for different acid treatments, all steps were conducted under identical conditions.

### Chemiluminescent detection

All blots were detected using a regular ECL method. The ECL solution A and B were mixed in a ratio of 1:1 to form a working solution prior to each detection. Then, the blots were treated with the working solution for 1 min at RT. After draining off excess working solution, the blots were either imaged with the QinXiang ChemiScope 6200 Touch (PRC) system or visualized with the traditional X-ray film method (Beyotime, PRC).

### Densitometric quantification

Densitometric quantification of western blot band intensity was performed by using the ImageJ software. The band intensity of each protein was first normalized to the respective loading control GAPDH. Relative expression level was further calculated as a ratio of the young group (DN101).

### Statistical analysis

Densitometric data were presented as mean ± SEM. A two-tailed unpaired student’s t-test was performed using the GraphPad Prism (Ver. 8) software. Statistically significant differences between groups were indicated by **p < 0.01 and ***p < 0.001.

## RESULTS

### Efficient inactivation of HRP by PTSA

First, we performed a dot blotting analysis to evaluate the effect of PTSA on HRP activity. As shown in Figure 1A, the activity of HRP diluted from 1000 to 10,000-fold could be sensitively detected by a regular ECL method before PTSA treatment. The blots were then treated with PTSA for 5 min at RT. Surprisingly, when treated with 0.05-0.5 M PTSA, no any signal of the spots, even with the highest HRP concentration, was detected using the same ECL method. At the concentration below 0.05 M, PTSA also showed a dose-dependent inactivation of HRP. These results indicated that a brief treatment with PTSA from 0.05 M to 0.5 M for 5 min at RT could inactivate HRP quickly and efficiently (Figure 1B). Compared to the non-treated blot (PBST), 10% AA treatment, as a positive control, only caused a clear but insufficient inactivation of a wide concentration range of HRP (1:1,000∼1:8,000). In sharp contrast to that most AA-treated spots on the blots still remained very clear signal, the twin blots treated with PTSA showed no any signal of HRP activity, even on the spot with the highest concentration of HRP, when treated for only 1 min (Figure 1C). Therefore, this result showed that PTSA impressively outperformed AA for inactivating HRP.

**Figure 1.**
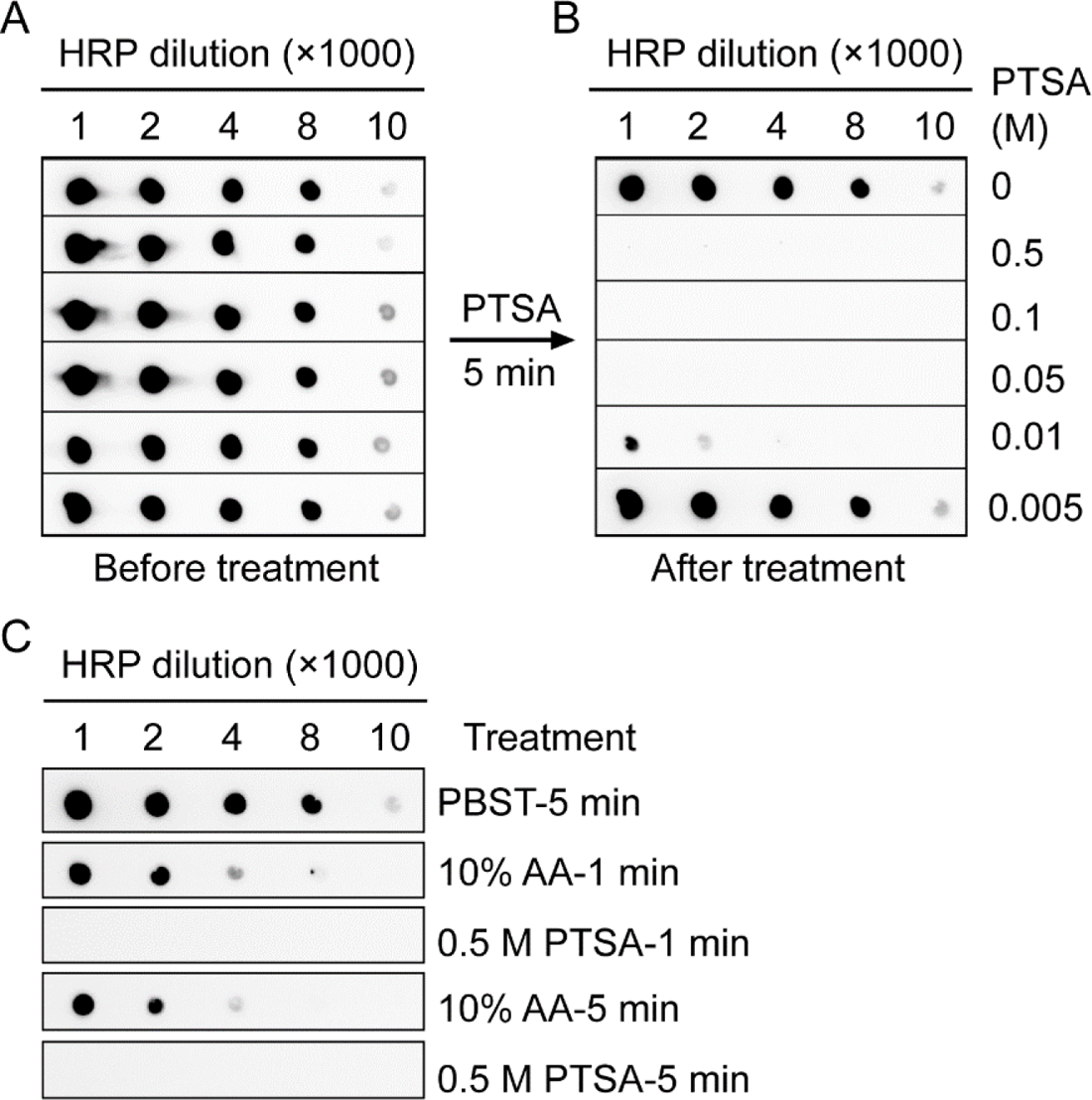
Dose-dependent inactivation of HRP by a brief PTSA treatment. A. Initial HRP activity before PTSA treatment. The activity of HRP as low as a 10,000-fold dilution could be sensitively detected by a regular ECL method. B. PTSA dose-dependently inactivate HRP. Note that no any signal of HRP was detected after a brief treatment (5 min, RT) with PTSA through a broad concentration range of 0.05-0.5 M. C. PTSA impressively outperformed AA for inactivating HRP. Dot blots with a wide concentration range of HRP remained clear signal after treated with AA for either 1 min or 5 min. No any signal of all the tested concentrations of HRP appeared on the blots treated with PTSA for only 1 min.

### Irreversible inactivation of HRP by PTSA without protein loss

To determine whether inactivation of HRP by PTSA is irreversible, several NC membranes were equally spotted with HRP-conjugated donkey anti-mouse IgG (1:1000), and then treated with PBST, 10% AA, and PTSA, respectively, for 1 hr at RT. Following treatment, the blots were thoroughly washed for three times in PBST to remove any residue acids, immersed in fresh PBST for 96 hr at 4℃, and then exposed with fresh ECL working solution. As expected, no any visible signal appeared on the 0.5 M PTSA-treated blots (Figure 2A), indicating that PTSA irreversibly deactivated HRP, while the major HRP activity appeared again on the blot treated with 10% AA (Figure 2A). Furthermore, the treated blots were stained with Ponceau S to assess the potential protein loss caused by each treatment. The intensity and size of all the Ponceau S-stained spots were nearly identical with and without PTSA treatment (Figure 2B), suggesting that there was no obvious protein loss after PTSA treatment.

**Figure 2.**
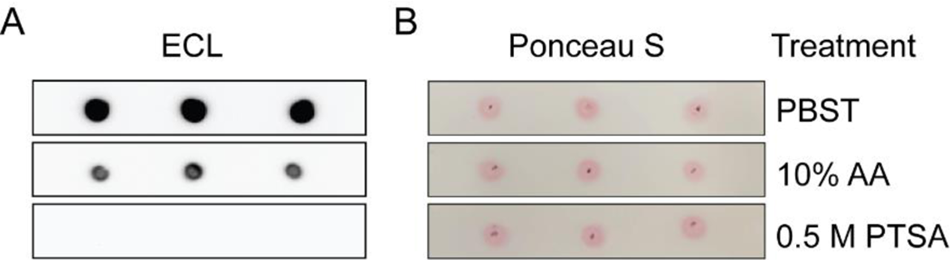
PTSA irreversibly deactivated HRP without protein loss on dot blots using NC membrane. A. Dot blots post-PTSA treatment showed no recoverable HRP activity after long term recovery storage, while blots parallelly treated with AA still generated clear signal. B. Similar protein levels were detected via a Ponceau S staining for the dot blots post-treatment with PBST, AA, or PTSA. Note that all NC membrane blots post-PTSA treatment kept neat and flat.

### Negligible epitope damage of PTSA-treated blots

HRP inactivation by PTSA on PVDF membrane was further evaluated after WB analysis. Protein lysates (10 μg per lane) from human fibroblasts were loaded in sextuplicate for SDS-PAGE and then transferred onto a PVDF membrane, followed by general immunodetection. One of the most abundant proteins, actin, was first detected with a mouse anti-actin antibody. The sextuplicate lanes on the blot showed single bands with equivalent signal of β-actin (Figure 3A). The membrane was then cut into 6 equal parts for parallel treatment. Each single part was separately treated with excess PTSA solution from 0.005 M to 0.5 M. After treated for 1 min at RT with constant shaking, they were thoroughly washed with PBST to remove any residue PTSA, and then immersed in fresh PBST for recovery at 4℃ for 96 hr. Further, the second protein, glyceraldehyde-3-phosphate dehydrogenase (GAPDH), was detected with a rabbit anti-GAPDH antibody, followed by the HRP-conjugated donkey anti-rabbit IgG antibody. Similar to the above dot blotting result, there was no any recovery of HRP activity for the band of β-actin throughout the most effective concentration range of PTSA (0.05-0.5 M) (Figure 3B). Once again, this result suggested that PTSA-mediated HRP inactivation is irreversible. Importantly, the band signals of GAPDH were detected equivalently, without observable difference, among all 6 blot parts treated with a broad concentration range of PTSA (Figure 3B). This strongly indicated that there was no visible sign of protein loss and epitope damage caused by PTSA treatment.

**Figure 3.**
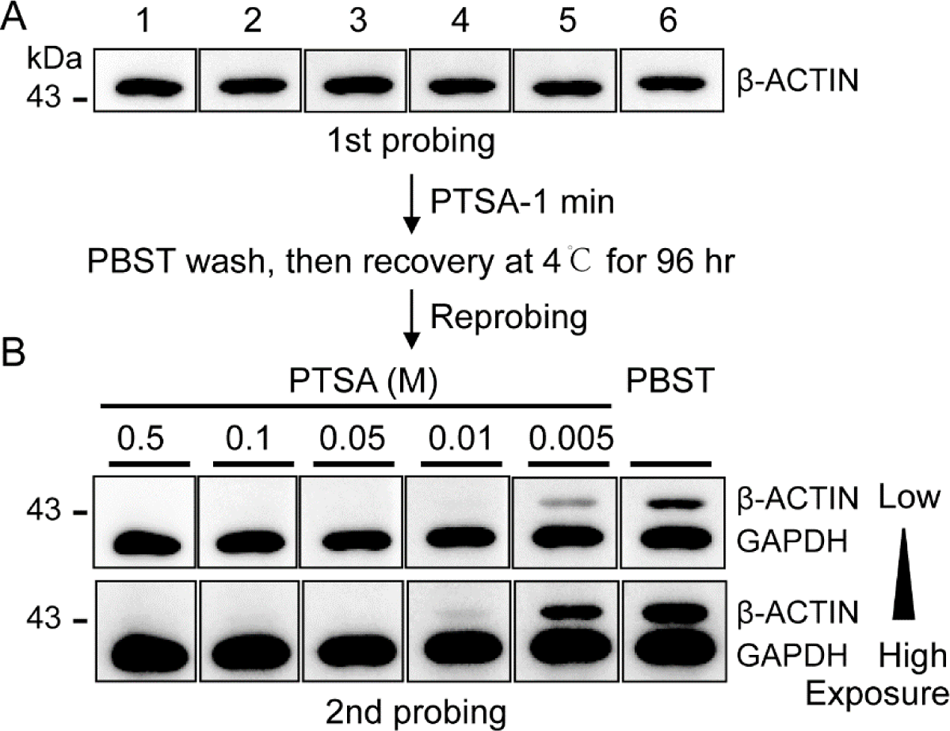
PTSA eliminated HRP activity without protein loss and epitope damage on western blots using PVDF membrane. A. All the sextuplicate lanes on the blot loaded with 10 µg/lane of fibroblast lysate showed single bands with equivalent signal of β-ACTIN. B. Efficient and irreversible inactivation of HRP by PTSA (0.05-0.5 M) permitted subsequential detection of GAPDH on the same blots. PTSA treatment showed a dose-dependent pattern in minimizing the interference from the residue signal of β-ACTIN. No protein loss and epitope damage of GAPDH appeared throughout a broad concentration range of PTSA and the treated PVDF membrane blots also kept neat and flat.

### Mutiple reprobing of PTSA-treated blots for human induced BFCNs

Since efficient HRP inactivation could enable multiple reprobing for a single blot, PTSA was applied for sequential detection of several proteins in small and precious sample. Neuronal lysates from a small-scale human induced BFCN have relatively low protein concentration, so as little as 1 µg per lane of total protein was loaded for SDS-PAGE and then transferred onto PVDF membrane. The same blot of BFCN sample was multiply probed with different antibodies to detect a total of at least 4 different proteins, including the neuronal markers class III beta-tubulin (TUJ1) and choline acetyltransferase (ChAT), the house-keeping protein GAPDH, and the potentially disease-relevant protein Galectin-3 (Gal3) [21]. Of note, a brief treatment with 0.05-0.5 M PTSA (1 min, RT) and a subsequent 5 min-wash with PBST were performed immediately before every next reprobing.

The less abundant cholinergic neuronal marker ChAT (∼68 kDa) was detected first (Figure 4A), followed by the abundant GAPDH (∼37 kDa) (Figure 4B), both using rabbit primary antibodies. This strategy successfully minimized the potential interference due to the cross-binding between the remained primary antibody for previous protein and the secondary antibody for the subsequent protein. Next, mouse primary antibodies were used to detect the third protein Gal3 (∼30 kDa) (Figure 4C), as well as the fourth protein TUJ1 (∼55 kDa) (Figure 4D). Overall, PTSA-mediated HRP inactivation, together with a proper strategy, such as probing per relative protein abundance and using antibody of different species, allowed fast and specific detection of multiple proteins, even for those with identical abundance or similar molecular weight, in the small and precious samples. In addition, because of no obvious protein loss and epitope damage post-PTSA treatment, there was no need of reblocking for the membrane. This further led to a significant reduction of the total time for reprobing multiple proteins, compared to the known HRP-inactivating or stripping methods.

**Figure 4.**
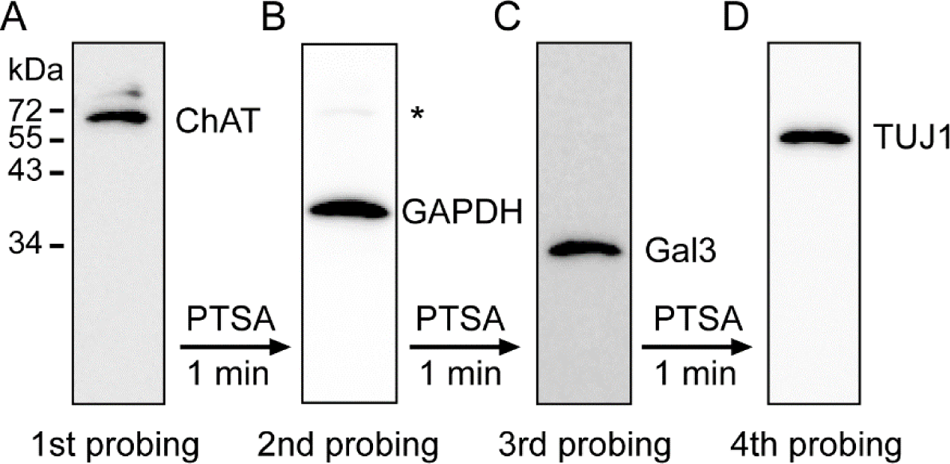
Inactivation of HRP by PTSA allowed sequential detection of at least 4 proteins on a single blot loaded with only 1 µg/lane of human induced BFCN lysate. A and B. Using respective rabbit primary antibody, the less abundant cholinergic neuronal marker ChAT was detected first (A), followed by detection of the house-keeping protein GAPDH (B). C and D. Subsequently, the disease-relevant protein Galectin-3 (C) and the general neuronal marker TUJ1 (D) were also detected on the same blot by sequentially using their corresponding mouse primary antibody. A brief PTSA-treatment (0.5 M, 1 min, RT) and following PBST-wash (5 min, RT) were conducted immediately prior to every next probing. No reblocking was needed after every PTSA treatment. Asterisk indicated a faint band of ChAT due to the cross-binding between the remaining rabbit anti-ChAT antibody and the donkey anti-rabbit secondary antibody for GAPDH.

### PTSA-mediated high-throughput WB analysis for 10 proteins of age-related dopaminergic hiNs

The above WB data from a single blot prompted us to develop a PTSA-mediated high-throughput WB method, in order to be able to detect diverse cohorts of proteins using only a single blot for young (DN101) and old (DN601) dopaminergic hiNs. Considering the highly restricted resource of such unique subtype of age-related neurons, only 1 µg per lane of each sample in triplicate was loaded for analysis. Following the strategies described above, up to 10 proteins were sequentially detected. Unless otherwise indicated elsewhere, the blot was treated with 0.05 M PTSA for only 1-5 min before every next probing and no re-blocking was conducted for all subsequent detection.

It has been reported that the age-/disease-related mitochondrial protein, peripheral-type benzodiazepine receptor/translocator protein (PBR/TSPO), mainly expressed in glial cells, was also expressed in dopaminergic neurons at much lower level [22-24]. Therefore, it was selected for the first probing. Consistent with its well-known role as an aging marker [24], the result showed a significantly up-regulated expression level of PBR/TSPO (∼19 kDa) in the aging dopaminergic hiNs (DN601) (Figure 5A and B). Similarly, LAMIN A/C (∼77/72 kDa), whose expression is known to be associated with age and disease [25], also showed an accumulation in the aging neurons (Figure 5A and B). Interestingly, another aging marker LAMIN B1 (∼73 kDa) [26] also obviously displayed an age-relevant expression pattern, with clearly declined protein level in DN601 (Figure 5A and B). Together, these comparable WB data from a single blot consistently validated the age-associate features of dopaminergic hiNs.

**Figure 5.**
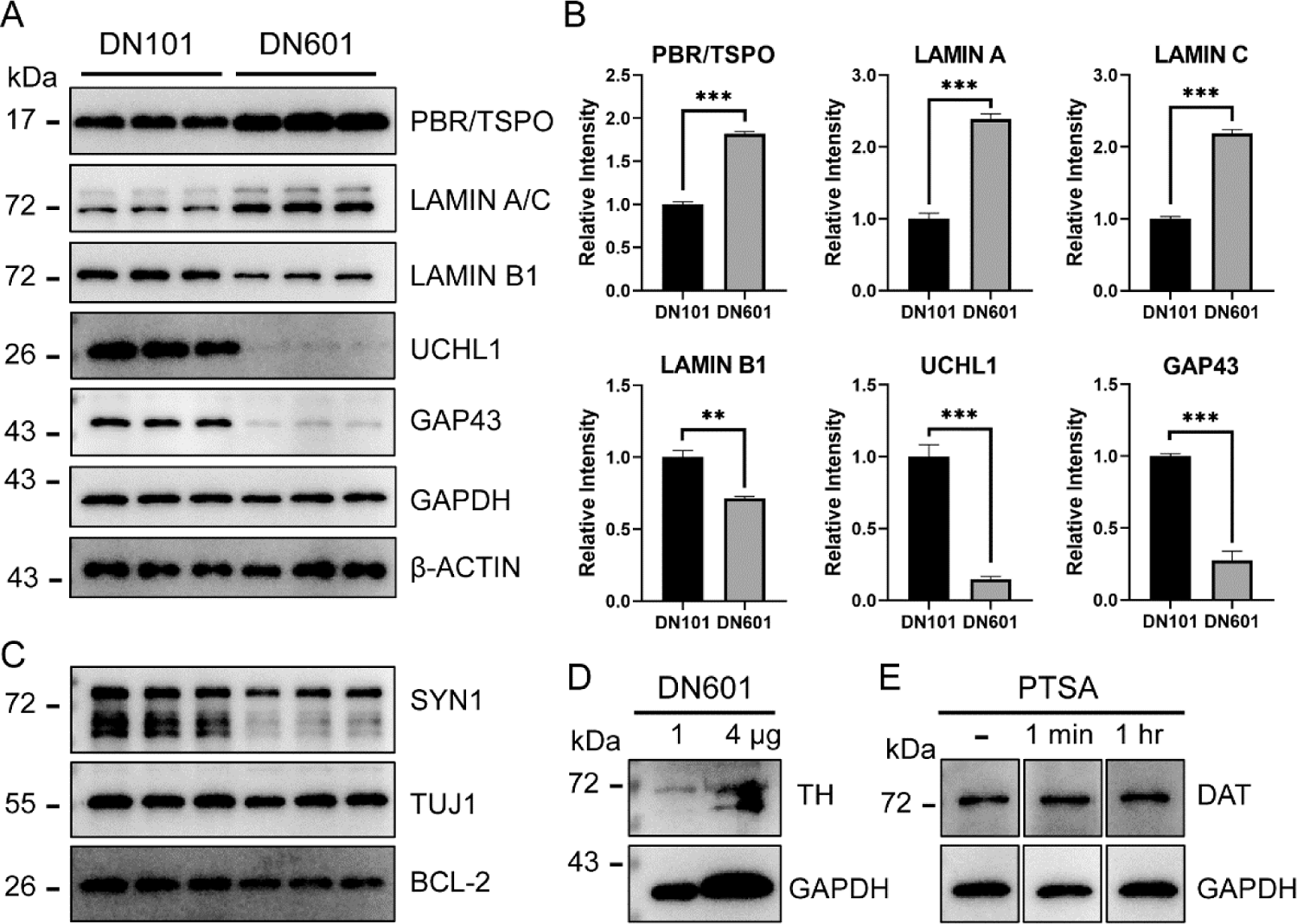
PTSA-mediated high efficiency western blot analysis for age-related subtype-specific dopaminergic hiNs. A. Typical expression patterns of three age-associated proteins (PBR/TSPO, LAMIN A/C, and LAMIN B1) and down-regulated expression of two PD-relevant proteins (UCHL1 and GAP43) in old neurons (DN601), compared to two loading controls GAPDH and β-ACTIN. Protein lysates of young and old dopaminergic hiNs were loaded for 1 µg per lane in triplicate. B. Quantification for the relative band intensity of aging- or disease-related proteins. Relative levels were calculated by first normalizing to the corresponding loading control GAPDH, followed by normalization to the level of young neuron (DN101). C. Two neuron-specific marker proteins SYN1 and TUJ1 were selectively detected almost without interference from previous proteins, and the neuroprotective BCL-2 was further detected by its specific antibody from different species. D. Dopaminergic neuronal marker TH was more clearly detected when the lysate of DN601 was loaded up to 4 µg per lane. E. Dopaminergic neuronal marker DAT was detected in the new blots (1 µg/lane) without or with PTSA treatment for 1 min or 1 hr. Quantitative data are presented as mean ± SEM. A two-tailed unpaired student’s t-test was performed using the GraphPad Prism (Ver. 8) software. **P<0.01 and ***P<0.001.

In addition, two PD-relevant neuron-specific proteins, ubiquitin C-terminal hydrolase L1 (UCHL1 or PGP9.5, ∼24 kDa) and growth associated protein 43 (GAP43, ∼43 kDa) [27-29], were also detected both in DN101 and DN601 (Figure 5A and B). Notably, the expression of UCHL1 and GAP43 exhibited a drastically aging-relevant reduction (Figure 5A and B), implicating them in a potential link with the late-onset PD. These results suggested a unique aging cell model and efficient WB analysis method for understanding mechanisms of late-onset neurodegeneration.

Meanwhile, several other proteins were also sensitively and specifically detected in this blot using their corresponding antibodies. These included the synaptic marker synapsin 1 (SYN1, ∼77 kDa), the general neuronal markers TUJ1 (∼55 kDa), and the house-keeping proteins GAPDH (∼37 kDa) and β-ACTIN (∼43 kDa) (Figure 5A and C). During the above high efficiency WB analysis, two pairs of proteins with an identical apparent molecular weight, including Lamin B1/Lamin C (∼72-73 kDa) and β-ACTIN/GAP43 (∼45 kDa), could be distinctly discerned by primary antibodies from different hosts (Figure 5A, Figure S1). In addition, using antibodies with high specificity, two or more proteins could be synchronously detected when they have a distinguishable size (Figure S1G, LAMIN B1 and β-ACTIN). Overall, the sensitivity did not appear to be affected post-PTSA treatment, as indicated by the detection of much less expression of UCHL1 and GAP43 in DN601 (Figure 5A).

Unexpectedly, the blot failed to be reused for probing the dopaminergic neuronal markers tyrosine hydroxylase (TH) and dopamine transporter (DAT). This might be due either to the poor sensitivity and specificity of certain antibody or to the potential interference from previously probed proteins, such as LAMIN A/C, LAMIN B1, and SYN1 (Figure 5A and C), because they showed a relatively overhigh abundance and a similar size range overlapping with TH and DAT, respectively. As an initial solution to this issue, we tried to increase the loading amount of sample. Indeed, when loading with 4 µg of lysate, two major bands of TH (∼57-72 kDa) could be more clearly detected using the specified antibody (Figure 5D and Figure S2A). Furthermore, a single band of DAT (∼72 kDa) was also selectively detected in the new blots loaded with only 1 µg per lane of the same sample (Figure 5E and Figure S2B). Not surprisingly, PTSA treatment for either 1 min or 1 hr did not impair the detection of DAT or GAPDH, but a brief 1 min-treatment before probing next protein GAPDH totally ruled out the interference from DAT (Figure 5E and Figure S2B).

Finally, we proved that this blot could still be used for detecting additional proteins of interest by using antibodies from different species. For example, a single band of the neuroprotective B-cell lymphoma-2 (BCL-2, ∼27 kDa) protein was successfully detected by its specific antibody derived from rats (Figure 5C). As a summary of the above results, all the original images, the antibody species, and the probing sequence of each protein were listed in the supplemental Figure S1A-J. During this PTSA-mediated high-throughput WB analysis, there were no observable protein loss and epitope damage for all the detected proteins. Taken together, we concluded that the relative protein abundance, the probing precedence, the primary antibody species, and the antibody sensitivity/specificity appeared to be key factors for successful detection of a variety of proteins by this PTSA-mediated high-throughput WB assay.

## DISCUSSION

WB analysis is one of the mostly cited methods for detecting specific proteins in a variety of samples throughout almost all fields of life science. Here, we described a modification to develop a high efficiency WB method for detecting cohorts of proteins on a single blot, in order to acquire more useful data with small and precious sample such as highly pure subtype-specific hiNs.

Similar to many previous efforts to solve this issue [15-18], we tried to develop an HRP-inactivating method, instead of stripping out the HRP-conjugated antibody. Meanwhile, we intended to avoid the use of currently known reagents, such as the chromogenic substrate, H_2_O_2_, NaN_3_, or AA, because of their respective drawbacks. Interestingly, we found that PTSA deactivated HRP on the dot or WB blots quickly and efficiently throughout a wide concentration range from 0.05 M to 0.5 M, and a feasible time-window from 1 min to 1 hr, even to overnight (data not shown). As far as we know, these results showed that PTSA drastically outperformed all the above mentioned HRP-inactivating reagents. In addition, PTSA has been used as a fixative for animal brain tissues that preserved with good morphology and histochemical properties [20], due to its unique feature as a strong, nonoxidizing organic acid. Consistently, we demonstrated that PTSA-treated protein blots kept neat and flat, without significant protein loss and epitope damage. Together, these unique capabilities of PTSA enable it to be used for immunodetection of multiple specific proteins on a single blot.

For successful multiple reprobing, it would be better to detect proteins of interest empirically from low to high abundance. Moreover, antibodies from different species were preferred for detecting proteins with similar abundance or close molecular weight. In combination with these strategies, we adopted a brief PTSA-treatment (0.05 M, 1 min at RT) and a subsequent rinse with PBST for 5 min before every next detection. We confirmed that this modification of the regular WB method made it much easier, faster, and more convenient than ever to detect at least 4 proteins on a single blot loaded with total protein as little as 1 µg per lane. Among them, the disease-related proteins Galectin-3 was also selectively detected in human induced BFCNs. Further research on the expression pattern of this inflammatory protein in BFCNs and other affected neuronal subtypes would be valuable for understanding the AD pathogenesis.

Many efforts for high-throughput WB assay have focused either on re-running gels and re-making blots quickly or on developing expensive and more advanced equipment [30-32]. However, new insights should be also considered, especially for small and precious samples with extremely limited availability. Multiple reprobing of a single blot provides a desirable solution for such an issue, in which the throughput is mainly dependent on how fast and efficient HRP can be inactivated without causing protein loss and epitope damage. Fulfilling with this requirement, our results identified that PTSA is an HRP-inactivating reagent well-suited for high-throughput WB analysis for several cohorts of proteins. Combined with a proper sequential strategy, simply repeating a brief PTSA treatment permitted multiple detection of up to 10 specific proteins on a single blot. Through such a flexible method, more informative data can be further obtained individually or synchronously from a single blot with highly specific antibodies. This led to not only a high efficiency analysis, but also a significant saving of time and precious protein samples.

Of note, our results from a single blot concurrently verified the expression of synaptic (SYN1) and general neuronal (TUJ1) proteins in dopaminergic hiNs. Moreover, the young (DN101) and old (DN601) dopaminergic hiNs respectively expressed the age-relevant proteins (PBR/TSPO, LAMIN A/C, and LAMIN B1) in a pattern in consistent with previous reports [22-26], further solidifying that hiNs retain age-associate features. Importantly, for the first time, two PD-relevant proteins, UCHL1 and GAP43 [27-29], were identified to be significantly down-regulated in the aging hiNs (DN601), bringing potential insights into the pathogenesis of late-onset PD. Together, these results pinpointed the great value of PTSA-mediated high-throughput WB analysis for small and precious samples, such as the highly pure aging-relevant subtype specific hiNs.

In conclusion, we demonstrated that PTSA inactivated HRP quickly and efficiently, allowing a simple, flexible, and high efficiency WB analysis of diverse proteins on a single blot for precious samples. It possesses the following advantages over known HRP-inactivating reagents: 1. Easy, fast, convenient, and efficient (0.05-0.5 M, 1 min, RT) for reprobing dot/WB blots; 2. No need of reblocking for next probing, further shortening the total time for multiple immunodetection; 3. No obvious protein loss or epitope damage, allowing multiple detection of specific proteins even with close molecular weight or abundance, when conducted with a proper strategy; 4. Non-volatile, colorless, no irritating odor, highly water soluble and stable, and pleasant to handle; 5. Easy to get and extremely low in price. As a strong organic acid, it should be noted that further investigation is needed to specify the compatibility of PTSA-treatment with certain specific proteins. In addition, it would be valuable to explore the versatile biochemical and immunochemical applications of PTSA following this study.

## AUTHOR CONTRIBUTIONS

Conceptualization, M.-J.L. and M.-L.L.; Methodology, Y.-F. S., M.-J.L., Z.L., X.S., and M.-L.L.; Investigation, Y.-F. S., M.-J.L., Z.L., and X.S.; Data analysis and interpretation, Y.-F. S., M.-J.L., Z.L., X.S., and M.-L.L.; Resources, Y.-F. S., M.-J.L., and M.-L.L.; Writing-draft, Y.-F. S., and M.-J.L.; Writing – Review & Editing, M.-L.L.; Supervision, M.-J.L., and M.-L.L. All authors reviewed and approved the final manuscript.

## ACKNOWLEDGMENTS

We thank Coriell Cell Repositories and ScienCell Research Laboratories for providing human fibroblasts.

## Supplementary Materials

**Figure S1.**
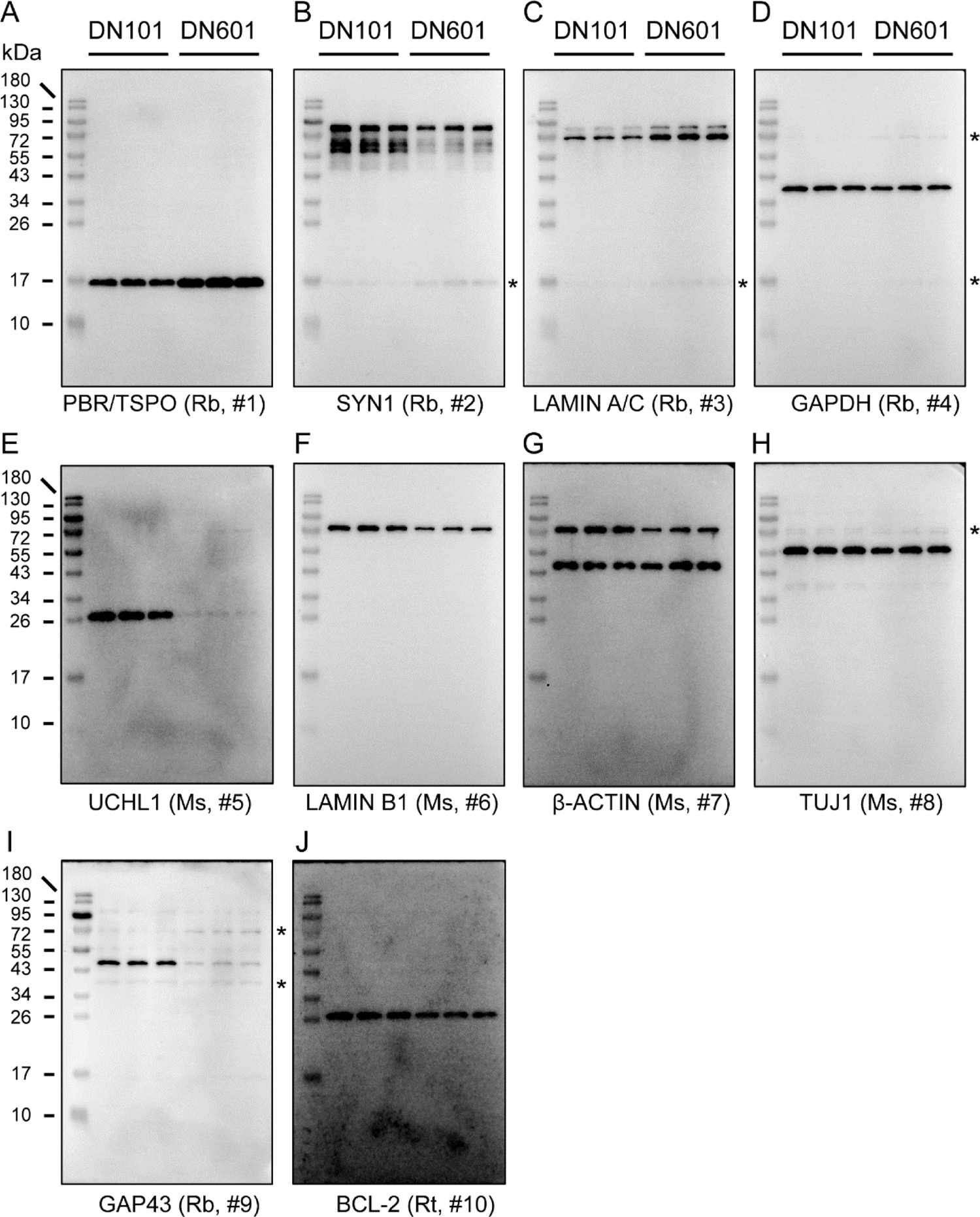
(Related to Figure 5A and C) Western blot original images and the probing sequence of 10 proteins. Rb, rabbit; Ms, mouse; Rt, rat. # Indicated the probing sequence of each protein. * Indicated some faint interference bands from previously detected high-abundance proteins when using antibodies from the same species. Such interference bands were negligible for detecting next protein with higher abundance and could be completely avoided when using antibodies from another species.

**Figure S2.**
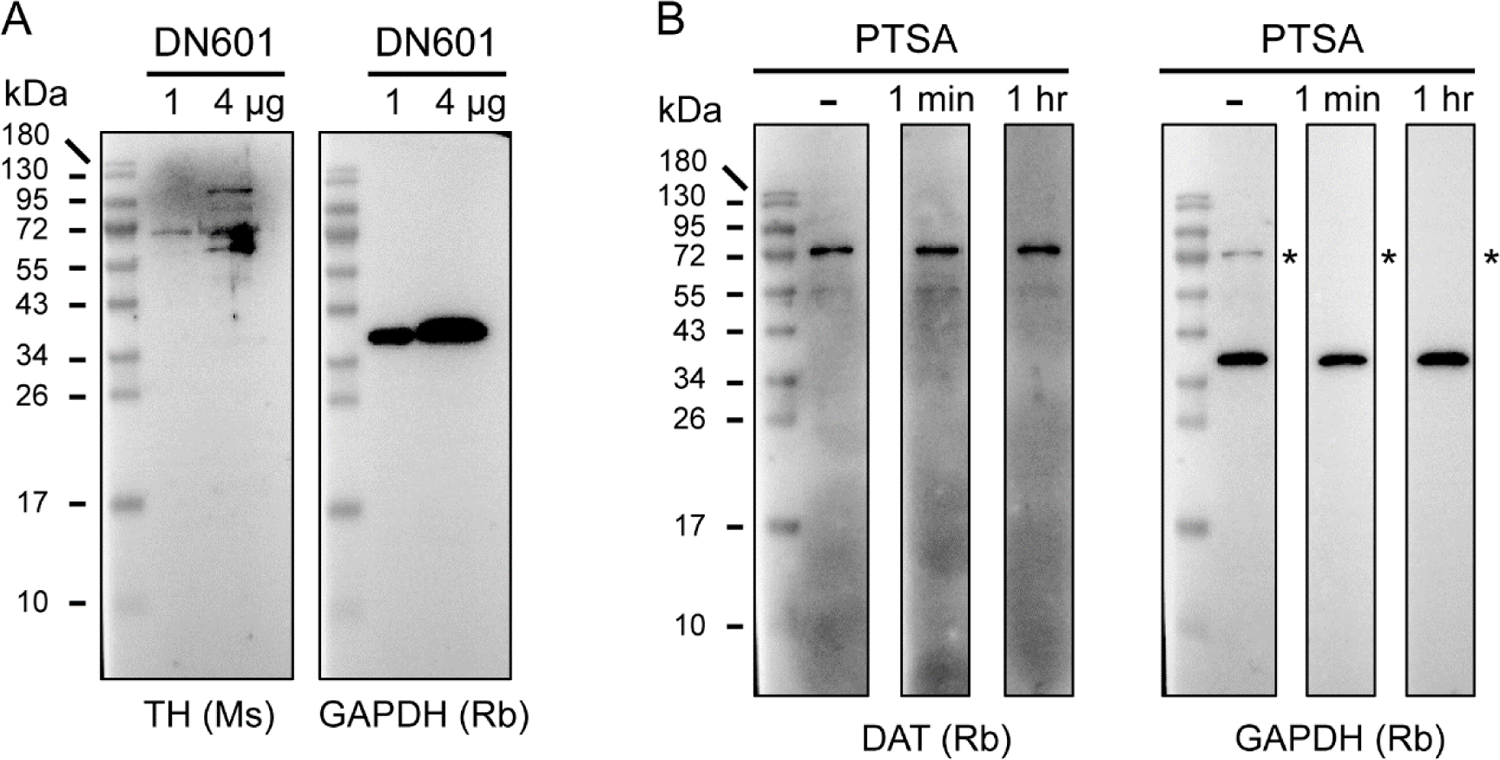
(Related to Figure 5D and E) Western blot original images of the dopaminergic neuronal markers TH and DAT. Ms, mouse; Rb, rabbit. * Indicated that, when detecting the house-keeping protein GAPDH with an antibody also from rabbit, the previously probed DAT band was still present in the blot without PTSA treatment, but was totally absent in the blots after PTSA treatment for 1 min or 1 hr.

